# Rapid changes in brain activity during learning of grapheme-phoneme associations in adults

**DOI:** 10.1101/2020.03.12.988402

**Authors:** Weiyong Xu, Orsolya Beatrix Kolozsvari, Robert Oostenveld, Jarmo Arvid Hämäläinen

## Abstract

Learning to associate written letters with speech sounds is crucial for the initial phase of acquiring reading skills. However, little is known about the cortical reorganization for supporting letter-speech sound learning, particularly the brain dynamics during the learning of grapheme-phoneme associations. In the present study, we trained 30 Finnish participants (mean age: 24.33 years, SD: 3.50 years) to associate novel foreign letters with familiar Finnish speech sounds on two consecutive days (first day ~ 50 minutes; second day ~ 25 minutes), while neural activity was measured using magnetoencephalography (MEG). Two sets of audiovisual stimuli were used for the training in which the grapheme-phoneme association in one set (Learnable) could be learned based on the different learning cues provided, but not in the other set (Control). The learning progress was tracked at a trial-by-trial basis and used to segment different learning stages for the MEG source analysis. The learning-related changes were examined by comparing the brain responses to Learnable and Control uni/multi-sensory stimuli, as well as the brain responses to learning cues at different learning stages over the two days. We found dynamic changes in brain responses related to multi-sensory processing when grapheme-phoneme associations were learned. Further, changes were observed in the brain responses to the novel letters during the learning process. We also found that some of these learning effects were observed only after memory consolidation the following day. Overall, the learning process modulated the activity in a large network of brain regions, including the superior temporal cortex and the dorsal (parietal) pathway. Most interestingly, middle- and inferior-temporal regions were engaged during multi-sensory memory encoding after the cross-modal relationship was extracted from the learning cues. Our findings highlight the brain dynamics and plasticity related to the learning of letter-speech sound associations and provide a more refined model of grapheme-phoneme learning in reading acquisition.

## 1. Introduction

The learning of grapheme-phoneme associations is a crucial step for reading acquisition in alphabetic languages. Unlike spoken language, written script is a recent cultural invention, and therefore no hard-wired brain circuit exists for reading at birth (Liberman 1992; Lieberman 2006). Consequently, reading acquisition involves plastic changes in pre-existing structural and functional networks of the brain, such as the visual and language systems to meet the new cognitive demand in reading (Dehaene et al. 2010; Dehaene et al. 2015). However, little is known about the cognitive processes and neural systems engaged during the learning of letter-sound correspondences, since most existing studies examine the long-term effects of learning to read.

One of the most important brain-level markers in learning to read is the increasing sensitivity to orthographic stimuli in the left ventral occipitotemporal cortex (vOT) (Brem et al. 2010; Ben-Shachar et al. 2011; Dehaene et al. 2010; Dehaene and Cohen 2011). The left vOT connects the visual word forms to other language areas of the brain, and has a posterior to anterior gradient (Lerma-Usabiaga et al. 2018; Vinckier et al. 2007), with the posterior part responsible for visual feature extraction and sensitive to smaller grain sizes (e.g., letters) and the anterior part to larger grain sizes (e.g., words (Dehaene et al. 2005)). Furthermore, vOT interacts with spoken language systems, for example, the phonological representations in the temporal cortex (Price and Devlin 2011). The development of letter specificity (greater activation for letters compared with false fonts) in left vOT is associated with reading ability in beginning readers (Centanni et al. 2018).

While changes in brain activity in the vOT is a hallmark for reading acquisition, learning to read requires other additional processes. A large body of neuroimaging studies (van Atteveldt et al. 2009; Raij et al. 2000; Blau et al. 2008) has looked into the audiovisual integration of graphemes and phonemes in literate adults who have mastered the associations through initial learning in childhood and years of the reading experience afterward. Consistent findings (Beauchamp et al. 2004; van Atteveldt et al. 2004; Richlan 2019; Blau et al. 2008; Calvert 2001; Wilson et al. 2018) suggest that the multisensory superior temporal cortex (STC) is the major brain region for audiovisual integration. For example, the brain activation to congruent audiovisual stimuli (letter-speech sound combinations) was found to be stronger than incongruent audiovisual stimuli in the left STC in transparent orthographies such as Dutch (van Atteveldt et al. 2004) and Finnish (Raij et al. 2000). In addition, the audiovisual congruency effect seems to be modulated by the transparency of the orthography as well as the task requirements used during the experiments (van Atteveldt et al. 2007). For example, audiovisual incongruent stimuli elicit larger neural responses than congruent stimuli in more opaque orthographies such as English (Holloway et al. 2015) and logographic scripts such as Chinese (Xu et al. 2019). The level of automaticity in audiovisual integration is important for normal reading development (Xu et al. 2018; Varga et al. 2018) and failing of which has been shown to be closely linked to dyslexia (Blomert 2011; Žarić et al. 2014; Blau et al. 2010). Automaticity for letter-speech correspondences seems to develop slowly, with electrophysiological mismatch negativity (MMN) studies showing a prolonged trajectory of audiovisual integration in children for up to 4 years after reading acquisition (Froyen et al. 2009). The slow development of automaticity is partly related to the neural representation of learned letters that must account for the numerous variations in position, case, and font for the same letter. This poses a great demand for reorganizing the hierarchical letter processing pathway along the vOT regions (Dehaene et al. 2005).

Brain processes for grapheme-phoneme associations have been studied for well-established associations in literate adults (Raij et al. 2000; van Atteveldt et al. 2004; van Atteveldt et al. 2007; van Atteveldt et al. 2009; Blau et al. 2008; Froyen et al. 2008) and children at different stages of learning to read (Žarić et al. 2014; Froyen et al. 2009; Froyen et al. 2011; Blau et al. 2010). These studies showed brain networks that are consistently activated during audiovisual integration days, months, or even years after learning of grapheme-phoneme associations. Less is known about the cognitive processes during the learning of new associations, which is arguably more complex and demanding than the automatic processing of existing associations. The scarcity of cross-modal studies on the learning process in humans is likely due to challenges in studying the brain mechanism during multisensory learning since it is very dynamic and involves multiple cognitive components such as sensory processing, multisensory integration, attention, memory formation, and consolidation.

The grapheme-phoneme learning process likely consists of several stages: First, during explicit learning, attention is directed to the information coming from the auditory and visual modalities. In addition, auditory and visual stimuli are combined into audiovisual objects in multisensory brain regions (Stein and Stanford 2008) (e.g., STC) and such cross-modal audiovisual association is initially stored in the short-term memory system. The short-term memories of audiovisual associations are consolidated during both practice and sleep (Diekelmann and Born 2010; Dudai 2012). They are then most likely transferred and stored in the neocortex for fast and automatic retrieval (Klinzing et al. 2019). Complementary learning systems have been shown in the medial-temporal systems (hippocampus and parahippocampal cortex) and neocortex, and a division of labor with the initial rapid learning in the hippocampus and gradual memory consolidation in the neocortical systems (McClelland et al. 1995; Davis et al. 2009). Converging evidence from a recent neuroimaging study also suggests a crucial role of the parahippocampal cortex for symbolic learning (Skeide et al. 2018). However, fast learning effects that occurred as a rapid form of memory consolidation at the time scale of seconds have also been reported in relation to motor-skill learning (Bönstrup et al. 2019). Such rapid consolidation might also play a role in other types of sensory learning (Hebscher et al. 2019).

Artificial grapheme-phoneme training paradigms that simulate the initial stage of learning to read in alphabetic scripts have provided interesting insights into the brain mechanisms of learning grapheme-phoneme associations. Learning-related brain changes have been reported at the time scale of minutes (Hämäläinen et al. 2019; Karipidis et al. 2017), hours (Taylor et al. 2014; Brem et al. 2018) and days (Taylor et al. 2017; Hashimoto and Sakai 2004; Madec et al. 2016; Quinn et al. 2017; Karipidis et al. 2018) after initial training. Combining information from visual and auditory modalities has been suggested to involve at least two possible mechanisms: Hashimoto and Sakai reported the involvement of the left posterior inferior temporal gyrus (PITG) and the left parieto-occipital cortex (PO) which showed plasticity for forming new links between orthography and phonology when learning novel letters (Hashimoto and Sakai 2004). The involvement of parietal brain regions has also been reported to be crucial for componential visual-verbal mappings in the early stages of learning to read (Taylor et al. 2014; Quinn et al. 2017). On the other hand, (Madec et al. 2016) have shown left vOT to be involved in phonological recoding processes of newly-learned letters by top-down influences from STG; this effect was affected by the strength of audiovisual associations in a two-day letter-speech sound training. Similar changes in the left vOT have been reported to show larger N170 responses (Brem et al. 2018) and vOT activation to trained than untrained characters after a short artificial character-speech sound training. These changes were also correlated with the training performance and were interpreted as a phonologically driven N170 and vOT tuning (Pleisch et al. 2019). Furthermore, these processes might be affected by modulation of attention to important features for learning in the frontal cortices (Hämäläinen et al. 2019). Interestingly the fast learning-related changes in brain activity seem to be linked to cognitive performance (Karipidis et al. 2018; Karipidis et al. 2017). Multisensory integration effects were found in a distributed brain network after a short letter-speech sound training (<30 min) in preschool children (Karipidis et al. 2017) with promising implications for identifying poor-reading children and predicting reading outcomes in pre-readers (Karipidis et al. 2018).

Despite the emerging insights from the available literature, to date, there is no comprehensive theoretical model of the cognitive processes and their brain level equivalents that are utilized during grapheme-phoneme learning. It is unclear when and how the audiovisual congruency effect starts to emerge in the multisensory superior temporal cortex and how quickly during training the visual representation of learned letters starts to differ from unfamiliar letters. Also, the allocation of attention is essential during explicit learning, yet how attentional processing is modulated by the learning material is still unknown. Finally, brain changes during the early stages of memory consolidation are still poorly understood, for example, effects related to repetition and practice during the initial learning stage, and the effect of overnight sleep on memory consolidation of letter-speech sound associations.

In our study, we investigated the neural mechanisms during the learning of novel grapheme-phoneme associations and the effect of overnight memory consolidation. The learning progress was tracked at a trial-by-trial basis during training on two consecutive days and was used to identify and segment different learning stages. Learning effects specifically related to grapheme-phoneme associations were studied by using two audiovisual stimulus sets. In one set, the audiovisual associations could be learned (cross-modal associative learning); in the other set, this was not possible due to the absence of information on the correct cross-modal correspondences (cross-modal non-associative learning). Different learning cues were presented after the audiovisual stimuli to dissociate the learning of correct audiovisual associations from basic multi-sensory processes. During cross-modal associative learning, the auditory and visual inputs had to be integrated and encoded into one audiovisual object, while no such integrative processes were needed in non-associative learning. We expected to see distinctive cognitive processes related to attention and memory encoding in non-associative and associative learning. Furthermore, we hypothesized that the learning of grapheme-phoneme associations would change the corresponding unisensory visual processing and elicit multisensory congruency effects. The unisensory effects were expected to occur in occipital and parietal regions due to the learning of the phonological representation of the Learnable letters mostly at a relatively late time window around 400 ms based on earlier studies (Xu et al. 2019; Xu et al. 2018; Quinn et al. 2017; Taylor et al. 2014; Dehaene et al. 2010). The multisensory congruency effects were expected to be elicited in the posterior superior temporal cortices in the late time window only after the learning of audiovisual associations (Wilson et al. 2018; van Atteveldt et al. 2004; Xu et al. 2019). These cross-modal learning effects were expected to be modulated by overnight memory consolidation. Finally, learning performance was correlated with cognitive skills linked to reading and working memory to explore the key behavioral factors that contribute to multisensory non-associative/associative learning speed.

## 2. Material and methods

### 2.1 Participants

In total, 36 people were scheduled to participate in the study. The participants were university students and staff recruited through email-lists and posters. The data from 6 participants were not included: 2 were excluded due to a low learning accuracy during the whole training session on Day 1, and for the other 4 participants, the MEG data were not measured because of cancellation. Therefore the data from the remaining thirty participants (20 females; 26 right-handed, 2 ambidextrous; mean age 24.3 years, SD 3.5 years, range 19-36 years) were used in this study. All participants had normal hearing and normal or corrected-to-normal vision (based on self-report). Participants were screened for the following exclusion criteria: head injuries, ADHD, neurological diseases, medication affecting the central nervous system, delays in language development, or any other language-related disorders. The ethical approval for this study was obtained from the Ethics Committee of the University of Jyväskylä, and the study was carried out in accordance with the Declaration of Helsinki. Participants gave their written informed consent prior to their participation in the experiments. After the MEG experiments, all of them received movie tickets or gift cards (for an equivalent value of 30 euros) as compensation for their time in the MEG recording and cognitive test sessions (see details of cognitive tests below).

### 2.2 Stimuli and task

Visual stimuli consisted of 12 Georgian letters 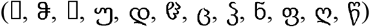. Auditory stimuli consisted of 12 Finnish phonemes ([a], [ä], [e], [t], [s], [k], [o], [ö], [i], [p], [v], [d]; mean duration: 473ms; SD:103ms). The auditory and visual stimuli were divided into two sets with 6 audiovisual pairs in each set. The stimuli between set one 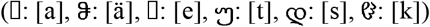 and set two 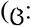 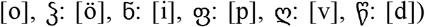 were chosen to match as closely as possible in visual and auditory complexity. The auditory stimuli in each of the Learnable and Control sets included three vowels and three consonants. The types of the phonemes were counterbalanced between the two sets (vowels were picked in the closed-open dimension: [e], [ä], [a] and [i], [ö], [o]; stop consonants: [p], [d], [t], [k]; fricative consonants: [s], [v]). The visual letters in the Learnable and Control sets were roughly matched in the overall shape and curvatures. Four additional letter-sound pairs 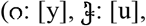 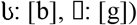 were used only for experimental instruction and practise purpose at the beginning of the experiment.

The audiovisual learning experiment consisted of 12 alternating training and testing blocks on Day 1, and 6 training and testing blocks on Day 2. In the training block, one of the two audiovisual stimulus sets was used as the Learnable set in which different learning cues (⍰ for congruent pairs (AVC) and X for incongruent pairs(AVI)) were presented after the simultaneous presentation of audiovisual stimuli. The other audiovisual stimuli set was used as the Control set, in which the feedback was always ▧ after the randomly-paired audiovisual stimuli (AVX). The audiovisual trial started with a fixation cross presented for 1000 ms, followed by the audiovisual stimuli for another 1000 ms. After the audiovisual stimuli, one of the three learning cues (“YES”: ⍰; “NO”: X; “UNKNOWN”: ▧) was presented for 1500 ms, depending on the types of audiovisual stimulus. Learnable and Control audiovisual stimuli were mixed and presented randomly in each training and testing block. In addition, auditory-only and visual-only stimuli from both sets were also presented in the training block, randomly mixed with the audiovisual stimuli.

Each training block was followed by a testing block in which the audiovisual stimuli from the training block were presented in random order, followed by a question on the stimulus congruency. The participants saw the question (“Match?”) displayed in the upper part of the screen and had to choose from three options (“YES”: ⍰; “NO”: X; “UNKNOWN”: ▧) provided on the lower part of the screen using a response pad. The order of the 3 options was randomized so that the participants would not learn to associate specific response buttons with specific options. After finishing the testing block, the feedback was provided about the accuracy of the previous block and all blocks they had done so far.

This was followed by a break in which participants were instructed to press a button to start the next training block when they were ready. The Learnable and Control sets were counterbalanced between the participants. Instructions for the participants and a short practice were given prior to the actual experiment on Day 1. Figure 1 shows the stimuli and experiment design of the study.

**Figure 1.**
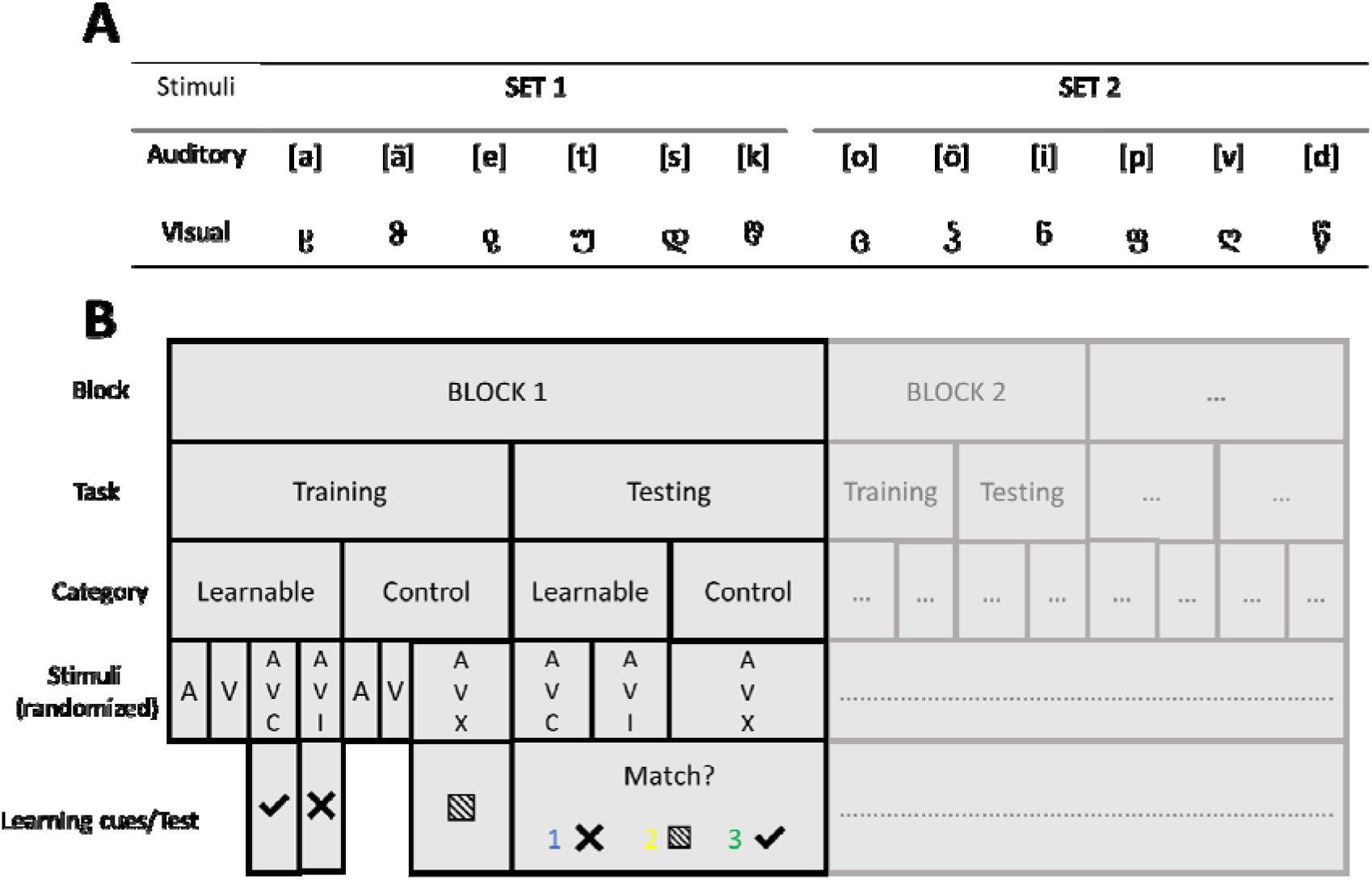
The experimental task on letter-speech sound association learning. A) Auditory stimuli consisted of 12 Finnish phonemes. Visual stimuli consisted of 12 Georgian letters. The auditory and visual stimuli were divided into two sets (counterbalanced between participants as Learnable or Control) with 6 audiovisual pairs in each set. B) The audiovisual learning experiment consisted of 12 alternating training and testing blocks on Day 1, and 6 training and testing blocks on Day 2. After presenting the audiovisual stimuli, different learning cues were provided in the training block. In the testing block, learning progress was tracked by asking questions on the audiovisual congruence, which the participants had to answer using a response pad.

### 2.3 MEG recording

MEG data were collected using the Elekta Neuromag® TRIUXTM system (Elekta AB, Stockholm, Sweden) in a magnetically shielded room at the University of Jyväskylä. A sampling rate of 1000 Hz and an online band-pass filter of 0.1-330 Hz were used in the data acquisition settings. The participant’s head position with respect to the MEG sensor arrays in the helmet was tracked continuously with five digitized head position indicator (HPI) coils. Three HPI coils were placed on the forehead and one behind each ear. The MEG head coordinate system was defined by three anatomic landmarks (nasion, left and right preauricular points). The anatomical landmarks, the position of the HPI coils, and the head shape (>100 points evenly distributed over the scalp) were digitized using a Polhemus Isotrak digital tracker system (Polhemus, Colchester, VT, United States) before the MEG experiment. The electrooculogram (EOG) was recorded with two electrodes attached diagonally slightly below the left and slightly above the right eye and one ground electrode attached to the collarbone. The MEG was recorded in a 68° upright gantry position with participants sitting comfortably on a chair. The same preparation and setup were used on Day 2.

### 2.4 Cognitive tests

A number of cognitive tests were administered to the participants to ensure they did not have language-related learning problems. Additional behavioral tests were conducted to run correlational analyses between learning speed in the MEG task and cognitive skill levels. The behavioral tests included the following: block design (visuospatial reasoning), vocabulary (expressive vocabulary), and digit span (forward and backward; working memory) from the Wechsler Adult Intelligence Scales (Wechsler 2008). In the block design test, the participants were shown how to arrange blocks with red and white colors to form a design, and they have to build the same design. In more difficult sections, the participants are only shown the design in a figure, and they have to build it. In the vocabulary test, the participants hear a word, and they have to describe the meaning of that word. In the digit span test, a series of numbers is said to the participant, and they have to repeat them either in a forward or backward order. The mean of the standardized scores in these tests was 10 and the standard deviation was 3.

Phonological awareness was tested using the Phonological processing task from NEPSY II (Korkman et al. 2007). In this task, the participant is asked to repeat a word and then to create a new word by leaving out a syllable or a phoneme, or by replacing one phoneme in the word with another phoneme.

Non-word repetition task from the Neuropsychological test battery (NEPSY; Korkman et al. 1998) was used to measure phonological processing and verbal short-term memory. The number of correct items out of 16 was used as the score.

Rapid automatized naming (Denckla and Rudel 1976), in which pictures of five common objects or five letters had to be named as quickly and as accurately as possible. The objects and letters were arranged in five rows, each containing 15 objects. The task was audio-recorded, and the time in seconds was calculated from the recording to be used in the analyses.

Three reading tests were included: word list reading using a standardized test of word list reading (Häyrinen et al. 1999), the number of correctly read words in 45 s was used as the score; non-word list reading based on Tests of Word Reading Efficiency (Torgesen et al. 1999), the number of correctly read non-words in 45 s was used as the score; pseudoword text reading (Eklund et al. 2015), number of correctly read words and total reading time were used as the scores. Writing to dictation was also assessed in which the participant heard 20 words and had to write them on a sheet of paper. The number of correctly written words was used as the score.

### 2.5 Data analysis

#### 2.5.1 Behavioral analysis

The cumulative learning index for each audiovisual stimulus was calculated based on the performance in the testing blocks. More specifically, for each auditory and visual components in the audiovisual stimuli, the learning index was defined as 0 if the participant pressed the wrong response button, and N (N=1,2,3 …) for the Nth time of successful learning (defined as correct response for both AVC and AVI for Learnable set, and correct response for AVX for Control set). The learning index for a specific audiovisual stimulus was the average learning index of the constituent auditory and visual components. The learning index was applied to the auditory only, visual only, and audiovisual stimuli in the same learning block prior to the testing block. For the incorrect button press after at least one successful learning, the learning index was defined as missing values (−1) and excluded from the analysis. These incorrect responses most likely reflect forgetting and lapses in attention to the task. The learning index gives a detailed quantification of the learning progress for each audiovisual stimulus; a learning index of N (greater than zero) indicates that successful learning happened on the Nth time when the audiovisual stimulus was presented.

The performance on the task (reaction time and accuracy) in the testing blocks for Learnable and Control stimuli were separately averaged by block (Figure 2A). In total, there were 12 (Block Index 1-12) blocks on Day 1, and 6 (Block Index 13-18) blocks on Day 2. In addition, the reaction time was also examined using the cumulative learning index (see the previous paragraph) to show the detailed learning progress (Figure 2B).

**Figure 2.**
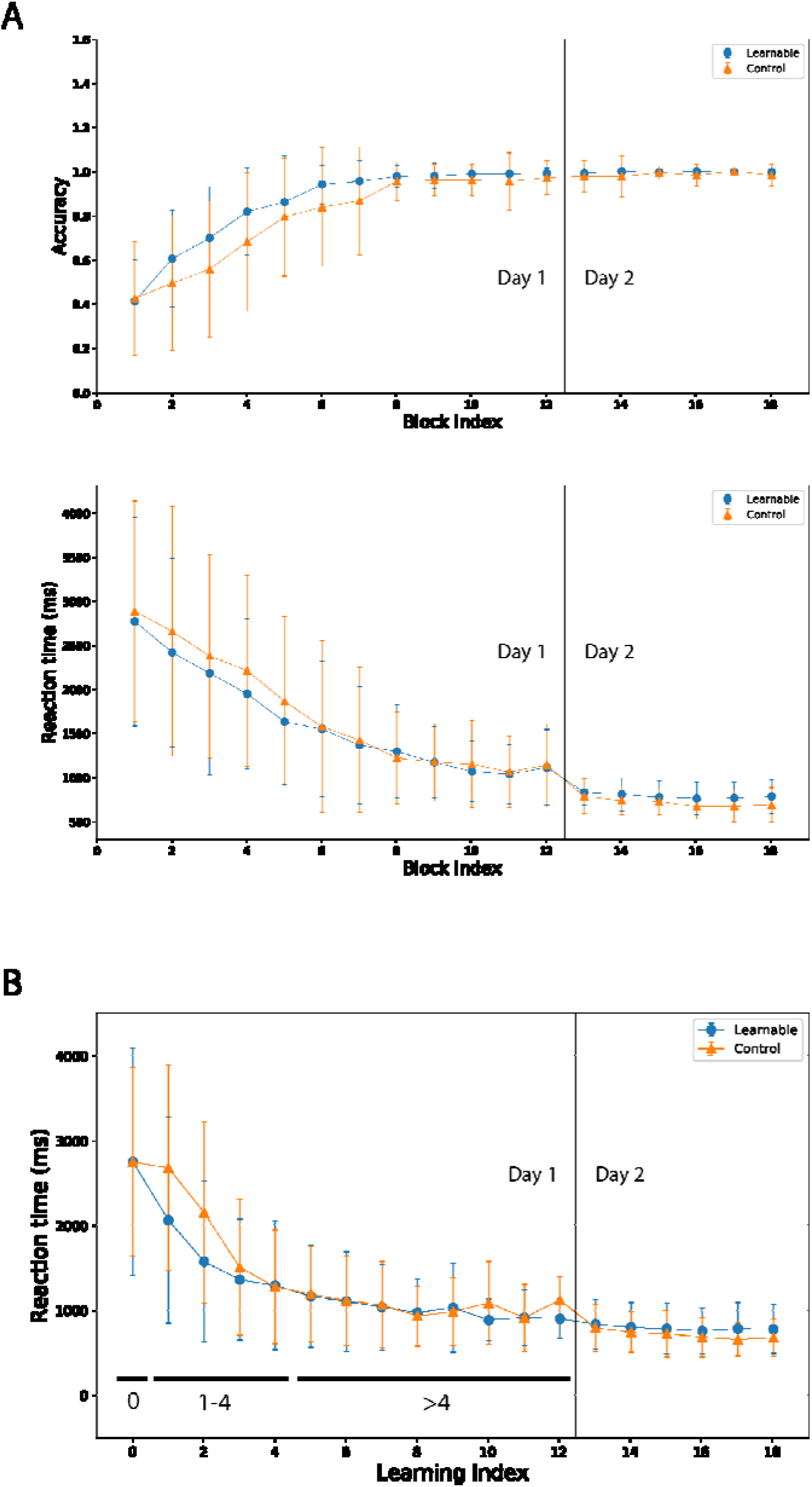
Task performance curve (mean and standard deviation) for Learnable and Control stimuli during the two-day audiovisual learning experiment averaged by block (A) and learning index (B). A) Reaction time and accuracy averaged by block. In total, there are 12 blocks (Block Index 1-12) on Day 1 and 6 blocks (Block Index 13-18) on Day 2 with a vertical line in the figure separating the two days. B) Reaction time averaged by the cumulative learning index. A learning Index of 0 indicates that the specific audiovisual pair has not been learned yet, and an integer number N greater than 0 indicates the Nth time of successful learning in the testing block for the audiovisual pairs.

Based on the learning progress, indicated by the reaction time (see Figure 2B), the participants acquire the letter-speech sound associations adequately after about 4 blocks of successful learning. The MEG data for Day 1 were therefore split over 3 learning stages (learning index=0,1-4 and >4) for the audiovisual conditions in learning and testing conditions separately. For Day 2, the MEG data were averaged together, since the participants had already learned all the audiovisual pairs. For the different learning cues, we postulate that the participants were paying attention to them before learning and immediately following the first few successful learning trials. This could be related to the fact that the short-term memory of the audiovisual stimuli is not well stabilized and consolidated, which could be seen from the fast decrease of reaction time when learning index=1-4 in Figure 2B. Therefore similar learning-related cognitive processes were presented when learning index=0-4 for the learning cue. To better capture the cognitive process in response to different learning cues, the MEG data were split into the following 3 parts for comparing the learning cues: learning index 0-4, learning index>4 on Day 1, and all the data on Day 2.

#### 2.5.2 MEG data analysis

MEG data were first pre-processed with Maxfilter 3.0.17 to remove external noise interference and compensate for head movement during the recording, using the movement compensated temporal signal-space separation (tSSS) method (Taulu and Simola 2006). Bad MEG channels were identified manually and were excluded; the time-series of the bad channels were reconstructed by Maxfilter.

Data were analyzed with MNE python (Gramfort et al. 2013) (version: 0.17.2). First, bad segments of MEG data were annotated and excluded from further analysis (Jas et al. 2018). MEG data were low-pass filtered at 40 Hz (zero-phase FIR filter design using the “hamming” window method). Fast ICA (Hyvärinen 1999) was then used to remove eye movement-related and cardiac artifacts. After applying ICA, data were segmented into epochs with 150 ms before and 1000 ms after stimulus onset. Bad epochs were first rejected based on peak-to-peak amplitude (grad=1500e-13 T/m, mag=5e-12 T) and then visually inspected in case of some remaining artifacts. Baseline correction was applied by subtracting the average response of the 150 ms prior to the stimulus onset from all data points throughout the epoch.

The multimodal interaction effects (i.e., processing of multi-modal stimuli is not merely the sum of auditory and visual stimuli presented separately) reflecting automatic and basic interaction between the auditory and visual processing were examined using the additive model (Audiovisual response = Auditory only response + Visual only response + Audiovisual interaction response). To calculate this regression analysis, the “linear_regression_raw” function in MNE Python was used. The interaction effects were calculated separately for the Learnable (LB) and Control (CT) stimuli by the linear regression analysis (AVC=A_LB_+V_LB_+Interaction_LB_; AVX=A_CT_+V_CT_+Interaction_CT_) for the three learning stages on Day 1 and Day 2. As can be seen from the model equations above, auditory response, visual response, and audiovisual interaction were the three predictors in the regression model. Since the MEG data was segmented into different learning stages, in order to get an adequate signal to noise ratio, a minimum number of 10 trials per average was used for including the MEG data in each regression analysis (Boudewyn et al. 2018; Luck 2005).

The fsaverage brain template from Freesurfer (RRID: SCR_001847, Martinos Center for Biomedical Imaging, Charlestown, MA, United States) was used for source reconstruction since individual MRIs were not available in the present study. Coregistration was done between the digitized head points and the template brain with a 3-parameter scaling. The average of the three scaling parameters was calculated for the recordings on Day 1 and Day 2 and was used for coregistration for both days to ensure the use of the same forward head model for each participant across the two days.

Depth-weighted (p = 0.8) minimum-norm estimates (wMNE) (Hämäläinen and Ilmoniemi 1994; Lin et al. 2006) were calculated for 10242 free-orientation sources per hemisphere. The dynamic statistical parametric maps (dSPM) (Dale et al. 2000) were used for noise normalization.

Region of interest analysis was used for comparing the estimated source activations to the three different audiovisual stimuli (AVC, AVI, AVX) in order to examine interaction effects in an analysis of variance (ANOVA) model. This was carried out for the training and testing blocks in 3 different learning stages on Day 1 and Day 2. Based on earlier literature (Karipidis et al. 2017; Xu et al. 2019; Raij et al. 2000) brain dSPM source waveforms (500ms to 800ms after stimulus onset) were extracted from left and right bank of the posterior superior temporal sulcus (labels: “bankssts”) (Calvert et al. 2001; Blomert 2011; van Atteveldt et al. 2009; Xu et al. 2019; Beauchamp et al. 2004; Wilson et al. 2018) as defined by the Desikan-Killiany Atlas (Desikan et al. 2006).

### 2.6 Statistical analyses

The audiovisual congruency effect was examined in a 3 (congruency: AVC, AVI, AVX) × 2 (hemisphere: left, right) repeated-measures analysis of variance (ANOVA) for the estimated activity in the region of interest analysis. To further examine possible audiovisual learning effects related to unisensory processing and multisensory interaction, spatiotemporal cluster-based permutation tests (Maris and Oostenveld 2007) were used for comparing Learnable and Control auditory, visual, and audiovisual interaction brain activations from the linear regression analysis based on the additive model.

Learning involves multiple cognitive components such as attention, active engagement, error feedback and memory formation and consolidation. In our experiment, these cognitive processes were reflected in the evoked responses elicited by the learning cues. To examine the cognitive processes that lead to successful learning of the AV associations, brain responses to different learning cues (“YES”: ⍰; “NO”: X; “UNKNOWN”: ▧) were also compared in pairs using the spatiotemporal cluster-based permutation tests. The number of permutations was set to 1000 for each test. The source data were downsampled to 200 Hz to reduce the computation time. The alpha threshold level was set to 0.05 for all tests.

Finally, to explore how much variance of the reading-related cognitive scores could be explained by the learning speed of Learnable and Control stimuli, correlation analysis (Pearson’s correlation coefficients) was carried out between the individual learning speed (average learning index of all Learnable and Control stimuli pairs in the twelfth block) on Day 1 and all the cognitive test scores (see the section above). The false discovery rate (FDR) was applied to correct the p-values in correlation analysis for the number of tests (Benjamini and Hochberg 1995).

## 3. Results

### 3.1 Cognitive skills and experiment performance

Descriptive statistics of the participants’ cognitive skill measures are presented in Table 1.

**Table 1.** Descriptive statistics of the participants’ cognitive skill measures (N=30). Standardized scores are shown within the parenthesis for block design, vocabulary, and digit span.

| Cognitive tests | Mean | SD | Range |
| --- | --- | --- | --- |
| Block design | 55.57 (12.1) | 9.97 (3.97) | 28 - 68 (2-19) |
| Vocabulary | 39.47 (13.03) | 7.49 (3.24) | 23 - 49 (6-18) |
| Digit span | 30.70 (12.43) | 4.61 (2.67) | 22 - 44 (7-19) |
| Nonword repetition, time (s) | 11.30 | 2.44 | 6 - 16 |
| NEPSY phonological processing | 51.80 | 1.03 | 49 - 53 |
| Rapid automatic naming(letters), time (s) | 17.93 | 4.48 | 10 - 29 |
| Rapid automatic naming(objects), time (s) | 31.73 | 4.49 | 23 - 40 |
| Word list reading, number of correct items | 104.77 | 0.50 | 103 - 105 |
| Word list reading, time (s) | 70.27 | 10.83 | 53 - 98 |
| Non-word list reading, number of correct items | 71.07 | 6.31 | 58 - 86 |
| Non-word text reading, time (s) | 29.97 | 5.35 | 22 - 42 |
| Non-word text reading, number of correct words | 37.20 | 1.10 | 35 - 38 |

As shown in Figure 2 A, the participants were able to learn the correct associations between the auditory and visual stimuli and to differentiate the Control stimuli from the Learnable stimuli. This was indicated by an increase in accuracy scores and a decrease in reaction times over training blocks. A similar decrease and stabilization of reaction times can also be observed by examining the reaction time averaged by learning index for Learnable and Control stimuli (Figure 2B). On Day 2, although the accuracy already reached ceiling level, there was a decrease of reaction time compared to the last block on Day 1 (Day 1 block 12: 1117 ms±436 ms v.s. Day 2 block 13: 825 ms±153 ms).

### 3.2 Grand average

The grand average of both sensor and source-level brain activities for auditory, visual, and audiovisual conditions on Day 1 and Day 2 are shown in Figure 3 (averaged across conditions). The activity patterns are typical for each stimulus type both in timing and in topography (localization). For example, the auditory responses were mainly localized around the perisylvian areas, and the visual responses were mainly localized in the occipital cortices.

**Figure 3.**
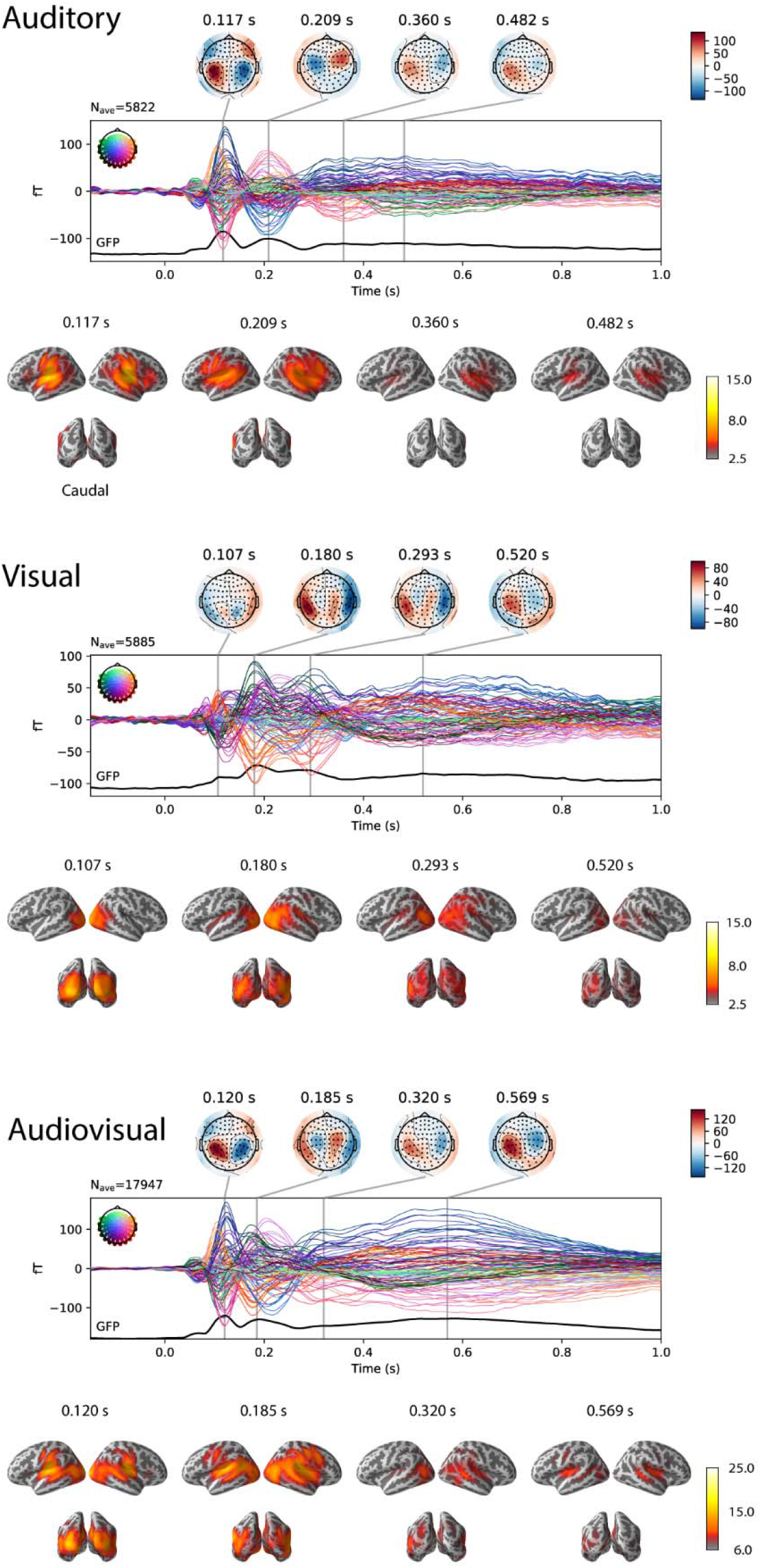
Grand average of both sensor and source brain activities for auditory, visual, and audiovisual conditions on Day 1 and Day 2. Top of each panel: Grand average of event-related field waveforms and topographic plots of major peaks defined by global field power from magnetometer channels. Bottom of each panel: Brain source dSPM activation at the major peaks as defined in the sensor level topographic plots.

### 3.3 Congruency effects in the STC

Repeated-measures ANOVA revealed significant main effects of the congruency on Day 1 only after learning of letter-speech sound associations in the training blocks (learning index >4: F(2, 52)=4.81, p=0.017) and in the testing blocks (learning index 1-4: F(2, 58)=4.37, p=0.022; learning index >4: F(2, 54)=4.43, p=0.022), as well as on Day 2 (F(2, 58)=3.82, p=0.034) during the training blocks.

Post-hoc t-tests indicated that dSPM activation to the Control audiovisual stimuli (AVX) was significantly lower (p<0.05) than to the Learnable audiovisual stimuli (AVC and AVI) in the Day 1 training blocks (when learning index: >4) and to the audiovisual congruent stimuli (AVC) in the Day 2 training blocks. During the testing blocks when the learning index was 1-4, the incongruent audiovisual stimuli (AVI) elicited significantly higher (p<0.05) activation than the Control audiovisual stimuli (AVX). The Learnable congruent audiovisual stimuli (AVC) elicited significantly higher (p<0.05) activation than the Learnable incongruent audiovisual stimuli (AVI) and the Control stimuli (AVX) in the Day 1 testing blocks when learning index was greater than 4. In addition, there was a hemisphere main effect (F(1, 29) =7.48, p=0.011) with higher dSPM activation in the right hemisphere than the left hemisphere during the training blocks on Day 1 at the stage when the learning index was 1-4. The results of the congruency effect are summarized in Figure 4.

**Figure 4.**
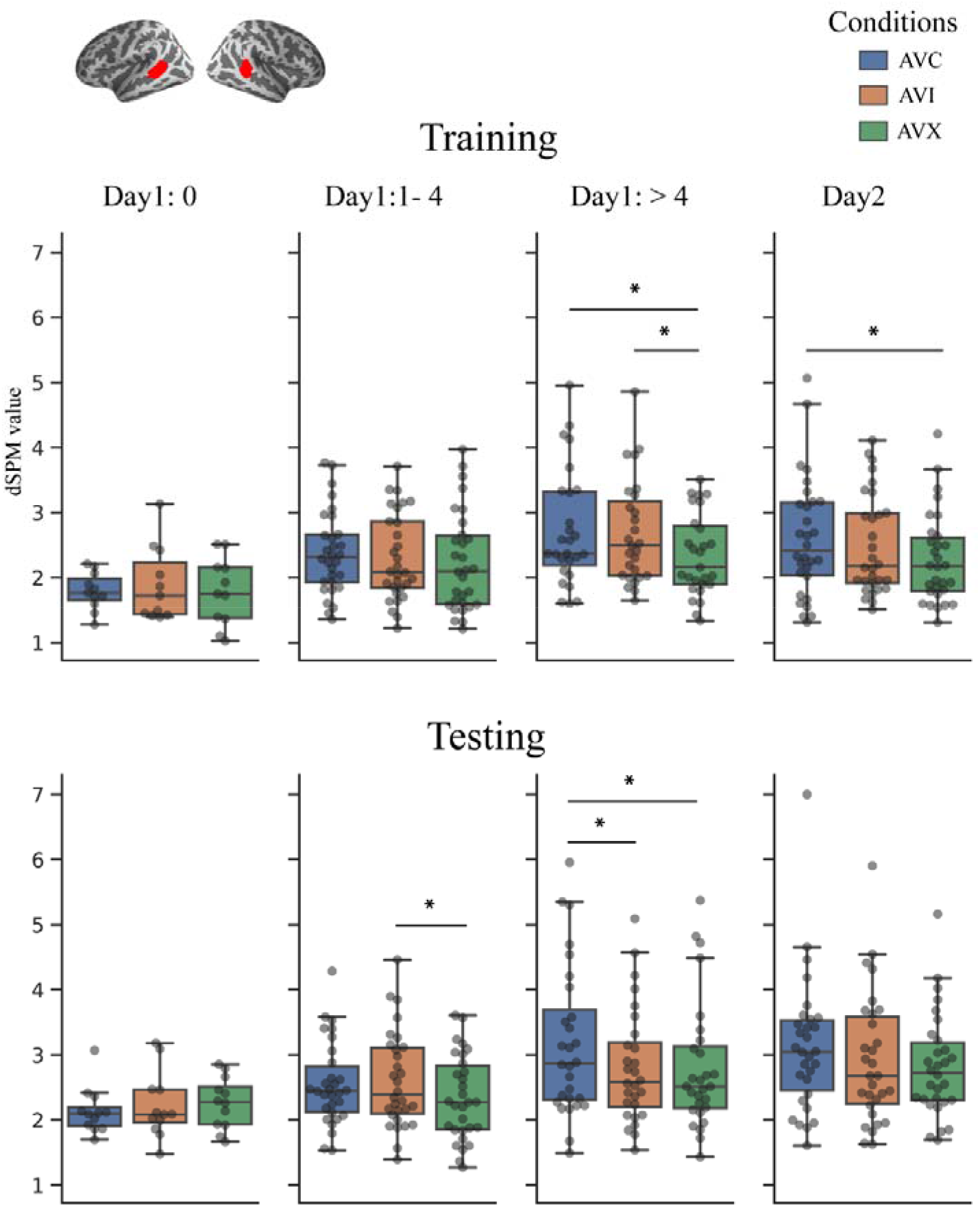
Congruency effect in the region of interest (left and right posterior superior temporal sulcus) for Day 1 and Day 2 during the training and testing blocks. The data were divided into three stages on Day 1 based on the learning indexes: 0, which means the letter-speech sound association has not been learned, learning index 1-4, and learning index >4. Since the participants had already learned all letter-speech sound pairs after Day 1, the data from all blocks on Day 2 were averaged. Results are shown in box plots with individual data marked as grey circles. Significant differences (p<0.05) are marked by horizontal bars and asterisks (*).

### 3.4 Auditory responses (Learnable vs. Control)

The learning effects were tested in the auditory modality by comparing the brain activations to the Learnable and Control auditory sounds from the time window of 100 ms to 800 ms after the stimulus onset using spatiotemporal cluster-based permutation statistics. The statistical tests were carried out for the 3 different learning stages (Learning index=0,1-4, >4 respectively) on Day 1 and the learned stage on Day 2. No significant differences were found for all the comparisons on Day 1 and Day 2.

### 3.5 Visual responses (Learnable vs. Control)

The learning effects were tested in the visual modality by comparing the brain activations of the Learnable and Control visual letters from the time window of 100 ms to 800 ms after the stimulus onset using spatiotemporal cluster-based permutation statistics. The statistical tests were carried between Learnable and Control stimuli for the three different learning stages (Learning index=0,1-4, >4 respectively) on Day 1 and the learned stage on Day 2. Significant differences were found between the Learnable and Control conditions on Day 1 when the learning index is greater than 4 (p=0.002, 455-795ms, left parietal, and occipital regions) and on Day 2 (p=0.001, 380-795ms, left parietal and occipital regions).

### 3.6 Audiovisual interaction effects (Learnable vs. Control)

The learning effects were tested for the audiovisual interaction by comparing the audiovisual interaction brain activations of the Learnable (AVC-A_LB_-V_LB_) and Control (AVX-A_CT_-V_CT_) conditions from the time window of 500 ms to 800 ms after the stimulus onset using spatiotemporal cluster-based permutation statistics. The statistical comparisons between the Learnable and Control stimuli for the three different learning stages (Learning index=0,1-4,>4 respectively) on Day 1 and the learned stage on Day 2. There was a significant difference (p=0.019, 500-680ms, left parietal region) at the stage when the learning index was 1-4 on Day 1.

Results for the auditory, visual, and audiovisual interaction comparisons between Learnable and Control conditions are shown in Figure 5.

**Figure 5.**
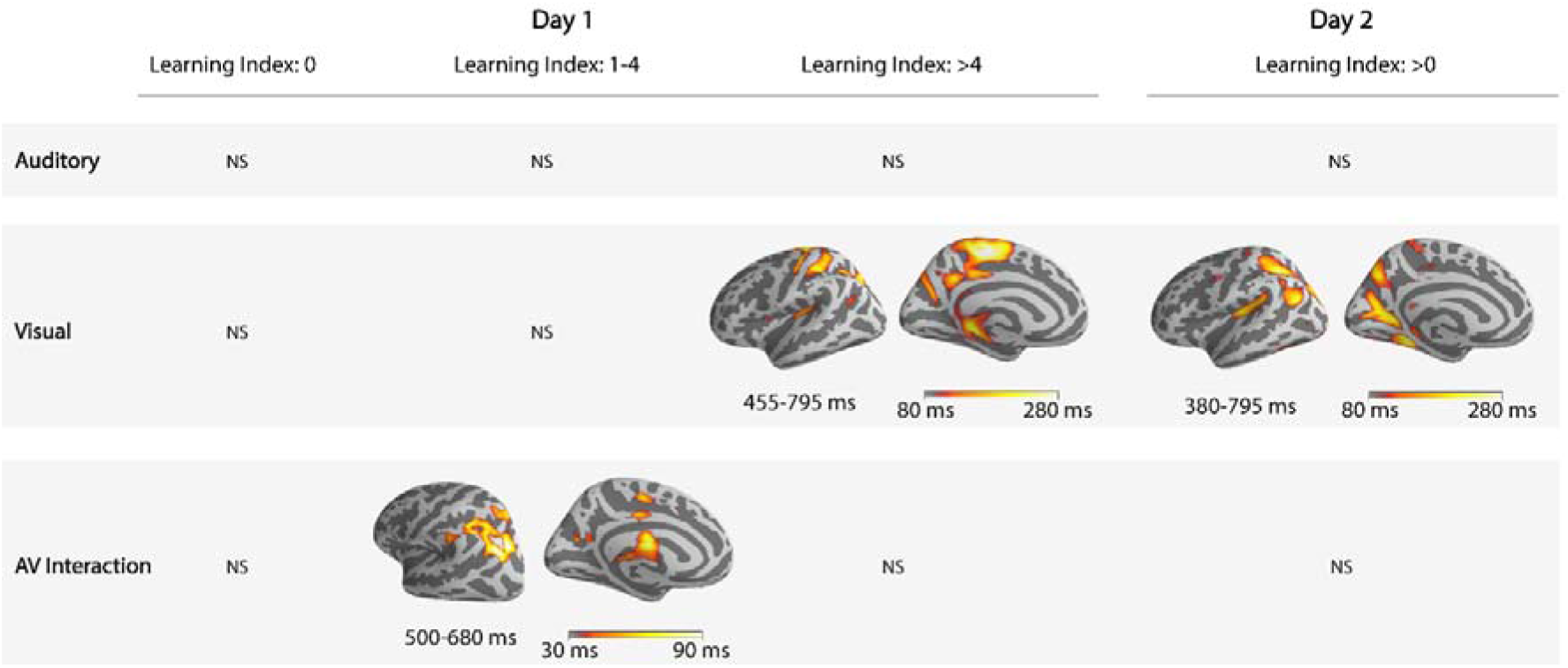
Learning effect for the auditory, visual, and audiovisual interaction conditions between the Learnable and Control stimuli on Day 1 (learning index: 0, 1-4, >4) and on Day 2 using spatiotemporal cluster-based permutation tests at the source level. The cluster on the basis of which the null-hypothesis was rejected is represented on the cortical surface, and the temporal duration of the cluster is indicated underneath the cortical surface. The brightness of the color on the cortical surface was scaled by the temporal duration of the cluster. Non-significant results are marked with NS.

### 3.7 Cortical responses to different learning cues

The cortical activities following the three different learning cues were compared in pairs using the spatiotemporal cluster-based permutation tests in the time window of 100 ms to 800 ms for Day 1 (Learning index=0-4, >4) and Day 2. There were significant differences between the 3 different learning cues when the learning index was between 0-4 on Day 1. Two clusters exceeding the threshold of randomization distribution under H0 were found for the ⍰ vs. X comparison, one (p=0.012) in the left temporal regions in the time window of 300-490 ms and another (p=0.016) in the right temporal regions in the time window of 295-550 ms. Two clusters were found for the ▧ vs. X comparison, one (p=0.008) in the left temporal regions in the time window of 360-730 ms and another (p=0.036) in the right temporal regions in the time window of 355-785 ms. Two clusters exceeding the randomization distribution under H0 were found for the ⍰ vs. ▧ comparison, one (p=0.040) in the left temporal regions in the time window of 400-780 ms and another (p=0.037) in the right temporal regions in the time window of 245-455 ms. In addition, there was a significant difference for the ▧ vs. X comparison (p=0.029, 300-740ms, left temporal, and occipital regions) when the learning index was greater than 4 on Day 1. No significant differences were found between the 3 different learning cues on Day 2. The results are shown in Figure 6.

**Figure 6.**
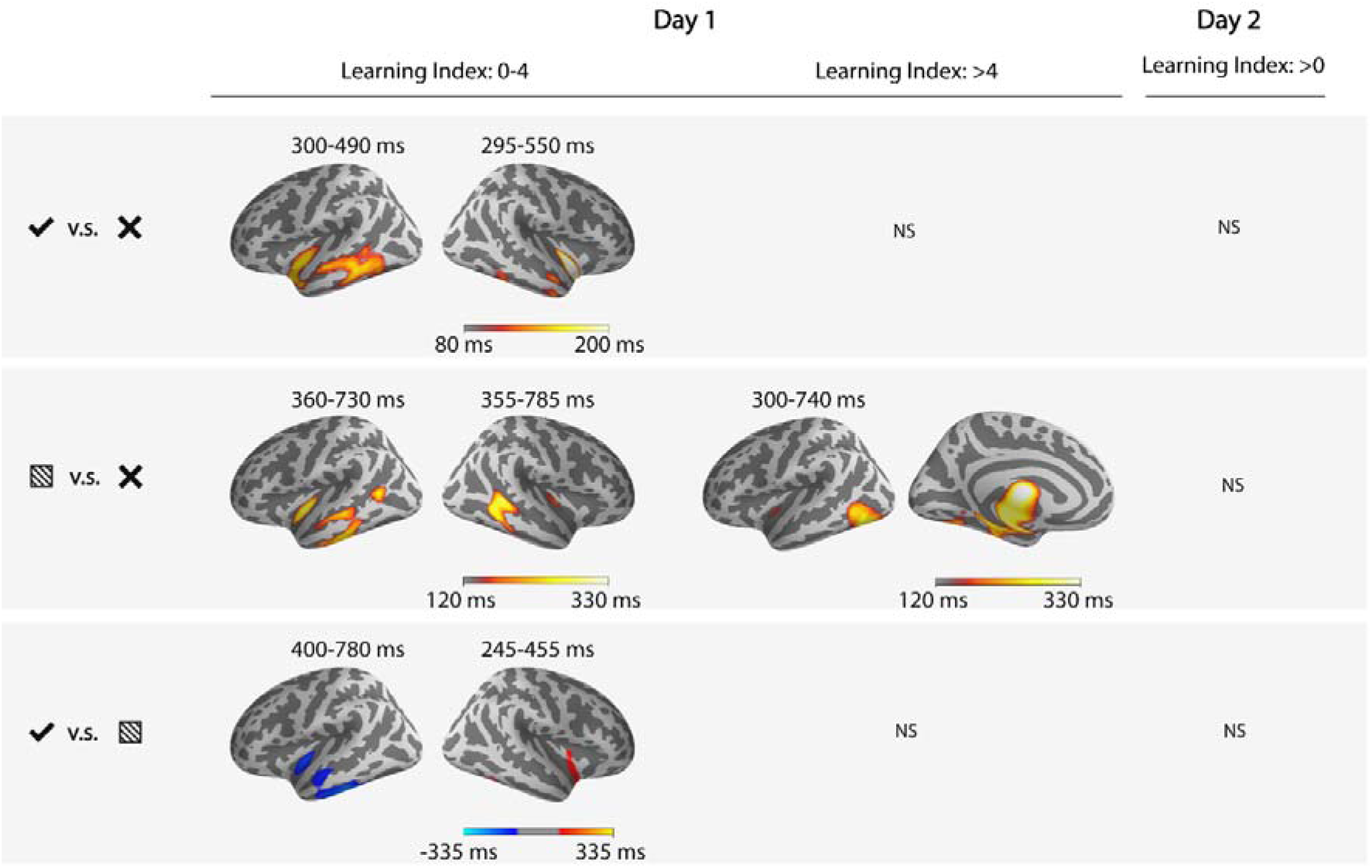
Comparison of cortical activity following the different learning cues at different learning stages. The activities following the three different learning cues were compared in pairs using the spatiotemporal cluster-based permutation tests in the time window of 100 ms to 800 ms. The MEG data were split into the following three parts: learning index 0-4, learning index>4 on Day 1, and all the data on Day 2. The time window of the cluster exceeding the threshold of randomization distribution under H0 is shown above each inflated brain. The colored region on the cortical surface is representing the cluster, and the brightness is scaled by its temporal duration. Warm color means the difference is greater than zero, and cold color means the difference is smaller than zero. Non-significant results are marked with NS.

### 3.8 Correlations between cognitive skills and learning speed

Correlation analysis was carried out between learning speed (of the Learnable and Control stimuli) and cognitive test scores. After FDR correction, only the learning speed of the Control stimuli was significantly correlated with the time spent on RAN objects (FDR-corrected p=0.000168). The results are shown in Figure 7.

**Figure 7.**
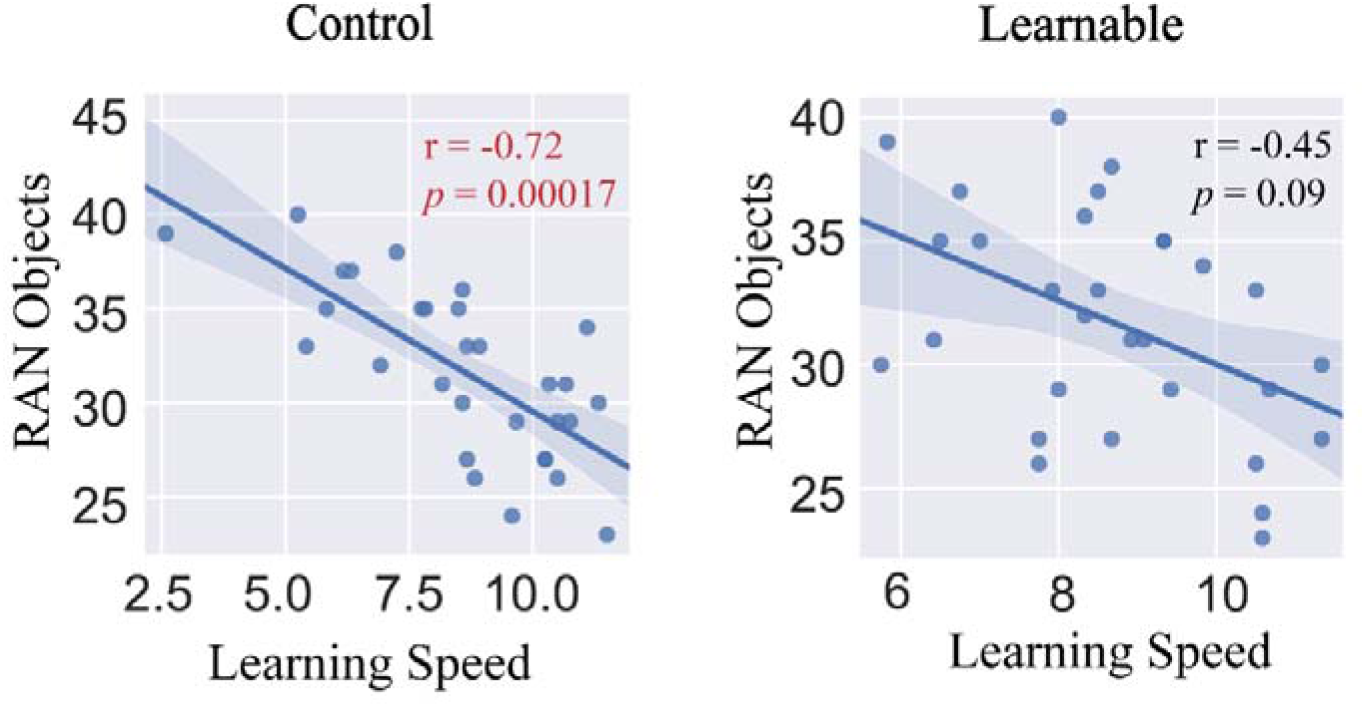
Correlations between learning speed and RAN objects. Significant correlations were marked with red color text, non-significant correlations in black.

## 4. Discussion

This study investigated the grapheme-phoneme association learning in adults. The cortical dynamics during initial learning and memory consolidation after learning were captured in a two-day letter-speech sound learning experiment using MEG. In the experiment, two sets of audiovisual stimuli were used for training in which the letter-speech sound association could be learned in one set (Learnable), but not in the other set (Control), based on the different learning cues provided. The experiment was designed to dissociate the audiovisual processing and the grapheme-phoneme associative learning by consecutive presentations of, first, the audiovisual stimuli, and second, different learning cues. The participants' performance was monitored with trial-by-trial precision in the testing blocks after each learning block. This allowed us to examine the changes related to associative learning by comparing the Learnable and Control conditions at different learning stages. These comparisons revealed dynamic changes in the brain processes during multisensory learning and, most interestingly, during the processing of the learning cues.

Region of interest analysis was conducted for comparing the brain responses to the audiovisual stimuli in the Learnable (AVC, AVI) and Control (AVX) sets in the posterior superior temporal sulcus (pSTS) in the time window of 500-800 ms based on earlier studies (van Atteveldt et al. 2004; van Atteveldt et al. 2009; Beauchamp et al. 2004; Raij et al. 2000; Karipidis et al. 2017; Xu et al. 2019; Wilson et al. 2018). As we expected, no significant differences were observed in the cortical responses to the different audiovisual stimuli before learning (learning index:0). The brain first started to process the audiovisual stimuli in the Learnable (AVI) and Control (AVX) sets differently in the early learning stages (learning index:1-4) in the testing blocks, and in the following training blocks (AVC>AVX and AVI>AVX) when learning index was >4 on Day 1. This suggested that the participants started to differentiate the Learnable and Control stimuli early in the learning process, which may reflect the easier differentiation of the two sets (Learnable and Control) compared to the learning of the audiovisual association within the Learnable set.

An effect of audiovisual congruency (AVC > AVI) in the left and right posterior superior temporal sulcus was found only at a later stage (learning index: >4) after the successful learning of letter-speech sound associations in the testing blocks on Day 1. This represents a brain level index of the learned associations. The stronger activation to congruent audiovisual stimuli than incongruent and control stimuli is in line with congruency effects reported in earlier studies using similar letter-speech sound learning paradigm (Karipidis et al. 2017; Karipidis et al. 2018). In addition, a similar effect has also been consistently reported for over-learned letters in literate adults (van Atteveldt et al. 2004; Raij et al. 2000) and has been interpreted as a result of language-related audiovisual functional connections that have developed during learning to read (van Atteveldt et al. 2004). However, this congruency effect in the testing blocks was absent on Day 2, possibly reflecting the effect of memory consolidation during sleep which could lead to the functional reorganization of multisensory memory (Rothschild 2019). Such functional reorganization of cross-modal connections might still be incomplete only one day after the initial learning, but with more practice and repetition could lead to the automation of letter-speech sound integration in literate adults (Froyen et al. 2009). The congruency effect has also been shown to be dependent on the experimental task (e.g., active/passive or explicit/implicit) (van Atteveldt et al. 2007; Blau et al. 2008). In our study, this is manifested as the different congruency effects for training and testing tasks at different learning stages in two days. For example, in contrast to the testing blocks, the brain responses to the Learnable audiovisual congruent (AVC) and Control (AVX) stimuli still showed a significant difference in the training blocks on Day 2. These differences most likely reflect the different cognitive processes during the training and testing blocks: e.g. active multisensory memory encoding was possibly mainly engaged during early training blocks, while memory retrieval and multisensory integration could only be possible after learning of audiovisual associations at later testing blocks. Therefore, these changes of congruency effects at different learning stages and during different tasks suggest dynamic characteristics of brain processes related to the newly-learned audiovisual associations.

Response to audiovisual congruence is not the only index reported in earlier literature that changes after learning the grapheme-phoneme associations. Therefore, more basic audiovisual interaction processes were also examined using the additive model (A+V vs. AV). The audiovisual interaction effect showed differences between the Learnable and Control conditions only at the early learning stage (learning index:1-4) on Day 1. Compared to the congruency effect, which is only possible after the crossmodal association has been learned (van Atteveldt et al. 2009), the (A+V vs. AV) comparison reflects more general form of cross-modal interaction, which has been shown to be important in children learning to read (Xu et al. 2018). The difference was maximal in the left parieto-occipital cortex, which has been indicated to be crucial for grapheme-phoneme mapping in learning to read (Sandak et al. 2004; Pugh et al. 2013; Bonte et al. 2017). Audiovisual interaction in the parieto-occipital cortex seems less often reported compared to the superior temporal cortex in many of the fMRI studies (Wilson et al. 2018; van Atteveldt et al. 2004; van Atteveldt et al. 2009). Earlier MEG studies using similar contrasts based on additive models (A+V vs. AV) have shown converging results in the parietal and occipital regions (Raij et al. 2000; Xu et al. 2018). Therefore this discrepancy could possibly be related to the different experimental paradigms and contrasts used in these studies to examine the audiovisual processes in learning to read. In the present study, the difference was only significant in the early learning stage, suggesting a transient learning process of actively combining auditory and visual information in the early learning stage for the Learnable set. In the later stages, after more stable multi-sensory memory representation was formed, no differences were found between the Learnable and Control stimuli.

Rather stable effects caused by learning of the audiovisual associations were found when examining the brain activation to the visual stimuli presented alone. The brain started to process the unimodally presented Learnable and Control letters differently at a later learning stage (learning index: >4), and this effect persisted on Day 2. The Learnable letters were closely linked to auditory stimuli (the phonemes) through repeated training on two days, whereas no such orthographic and phonological connection could be learned for the Control letters and phonemes. From the principle of Hebbian learning, seeing these Learnable letters alone should activate the phonemic representations of the letters. This seemed to occur after 4 repetitions of successful learning on Day 1 and continued to Day 2. Differences in phonological and orthographic processing of single letters vs. pseudo-letters have been reported (Bann and Herdman 2016; Herdman and Takai 2013) in both early (P1, N1, P2, and P3) and late (>300 ms) time windows using event-related potentials (ERPs). In our study, the time window of the cluster where responses to the Learnable and Control letters differed started from 455 ms on Day 1 and 380 ms on Day 2, which is relatively late compared to similar effects of learned letter vs. pseudo-letter comparisons in other studies (Herdman and Takai 2013; Brem et al. 2018; Maurer et al. 2005). This might reflect the very early stages of learning captured by the present study, which might exhibit a slower processing speed of grapheme-phoneme mapping than the processing of well-established or over-learned associations. The spatial extent of the clusters for the Learnable and Control letter comparisons showed widespread distribution around the left temporoparietal, paracentral, and occipital regions. The temporoparietal (dorsal) circuits including the angular gyrus and supramarginal gyrus (SMG) in the inferior parietal lobule have been associated with grapheme to phoneme conversion (Pugh et al. 2000; Sandak et al. 2004; Taylor et al. 2014; Bonte et al. 2017) and are suggested to show brain changes in early reading acquisition (Dehaene et al. 2015; Pugh et al. 2001; Carreiras et al. 2015). Corroborating evidence comes from a training study in which participants were taught to read artificial words (componential learning of letter-sound associations) and name artificial objects over two days with differences observed in the occipitotemporal and parietal regions when reading an artificial orthography compared to naming artificial objects (Quinn et al. 2017; Naumer et al. 2009).

With regard to unimodally presented auditory stimuli, we did not find any differences in the brain responses to the Learnable and Control sets. This was not surprising because the auditory stimuli were Finnish phonemes, which are familiar for native Finnish participants and are already closely linked to the Finnish letters through years of experience. Therefore, each phoneme is mapped to at least one Finnish letter and a newly learned Georgian letter for the Learnable set. However, mapping of additional visual information to existing phonemic representations might not alter brain representations of the existing phonemes. Instead, associations between the phonemic representation and a new visual representation would be formed, which is what we saw for the processing of audiovisual stimuli.

Overall, the results related to brain responses of the audiovisual stimuli suggested that multisensory processing is very dynamic and depends on the different learning stages and tasks. On the other hand, the effects on brain responses to the unimodally presented letters seem to be more persistent after successful learning. These early dynamic processes have not been reported before since most earlier studies have examined the multisensory or learning effects at one time point after training.

The findings described above are mostly brain-level indices of the learning results that occurred during the MEG recording. Of particular interest is, however, the brain mechanisms that lead to successful learning. Examining the brain responses to the three learning cues provided a unique window into brain processes that preceded the behavioral level of learning. Indeed, the brain responses to the learning cues were different mainly before and immediately after (learning index:0-4) behavioral learning could be observed on Day 1. On Day 2, after the audiovisual stimuli had been learned and consolidated, no differences were found between the brain activations to the three learning cues. In the training blocks, the auditory and visual information needs to be kept in working memory together with an initial weak association between the representations of the two different modalities after the presentation of audiovisual stimuli. The learning cue is processed first in the visual cortex (similar for all three cues) to extract the relevant information on the cross-modal relationship on the previously presented audiovisual stimuli, followed by the updating of that cross-modal relationship according to the cue. Both the associative learning process (reflected in the ⍰ vs. X contrast) and non-associative learning process (reflected in the ▧ vs. X contrast) showed differences in the left and right middle and inferior temporal and some deeper brain sources (labeled as insula in the Desikan-Killiany atlas (Desikan et al. 2006) probably reflecting short-term memory encoding of multisensory information. Differences found in the deep brain sources near the bilateral medial temporal regions, as well as the insula, could reflect the working memory processes in the hippocampus and related areas (Yonelinas 2013; Quak et al. 2015; Olson et al. 2006). Similar activation patterns have been reported in earlier studies; for example, the inferotemporal (IT) cortex has long been suggested to be important in forming associative long-term memory representations (Sakai and Miyashita 1991; Miyashita and Hayashi 2000) including audiovisual cross-modal associations (Gibson and Maunsell 1997). In addition, the occipitotemporal junction and parahippocampal gyrus have been reported to show increased activation when learning arbitrary cross-modal associations (Tanabe et al. 2005; Skeide et al. 2018).

The difference between associative and non-associative learning (⍰ vs. ▧) was mainly localized in parts of the left temporal and right insula regions. The decreased activation in left temporal regions might be related to the cross-modal memory encoding (Tanabe et al. 2005), and the increased activation in the right insula regions might be related to increased attention (Chen et al. 2015) for multisensory associative learning compared with non-associative learning. The relatively late time window of the effect (after about 300 ms) also suggests a multisensory working memory process engaged after the basic sensory processing of the learning cues. Therefore, cross-modal associative and non-associative learning utilized largely overlapping left and right middle and inferior temporal and deep brain regions (possibly reflecting activity from or activity modulated by, for example, insula or hippocampus) when using the ‘X’ cue as the baseline. However, these two types of learning also showed differential activation strength in both hemispheres, which probably reflects the unique cognitive processes in associative learning.

For the correlation analysis between learning speed and cognitive skills, we found that audiovisual non-associative learning speed was significantly correlated with the rapid naming of objects. Rapid naming ability, which is a robust behavioral precursor of reading fluency across various languages (Moll et al. 2014; Kirby et al. 2010), seems to be important for fast non-associative learning. This result is consistent with other studies using artificial language learning studies (Karipidis et al. 2017; Aravena et al. 2018; Karipidis et al. 2018), although this one correlation surviving the multiple comparison correction should be regarded with caution as the main variable of interest (the learning speed for the Learnable items) did not show significant correlations to the naming speed.

Based on findings from the present and previous studies, we formulate a global sketch regarding the learning of letter-speech sound associations in Figure 8. In this figure, the auditory (e.g., the sound of /a/) and visual (e.g., letter a) sensory inputs are first processed in the primary auditory and visual cortices. The auditory features of the stimuli are then combined to form more abstract representations most likely in the superior temporal regions in both early and late time window as reflected for example by the auditory P2 response (Hämäläinen et al. 2019) and late sustained responses (Ceponiene et al. 2005). The visual features have been combined and formed an abstract representation when the visual information is processed along the vOT cortex known to respond to orthographic regularities. The auditory and visual information are then integrated in the multisensory areas in the superior temporal cortex (van Atteveldt et al. 2009; Raij et al. 2000; Beauchamp et al. 2004) to form a coherent audiovisual object at a relatively late time window after the auditory and visual inputs are processed (see (van Atteveldt et al. 2009) for a functional neuro-anatomical model of letter-speech sound integration in literate adults).

**Figure 8.**
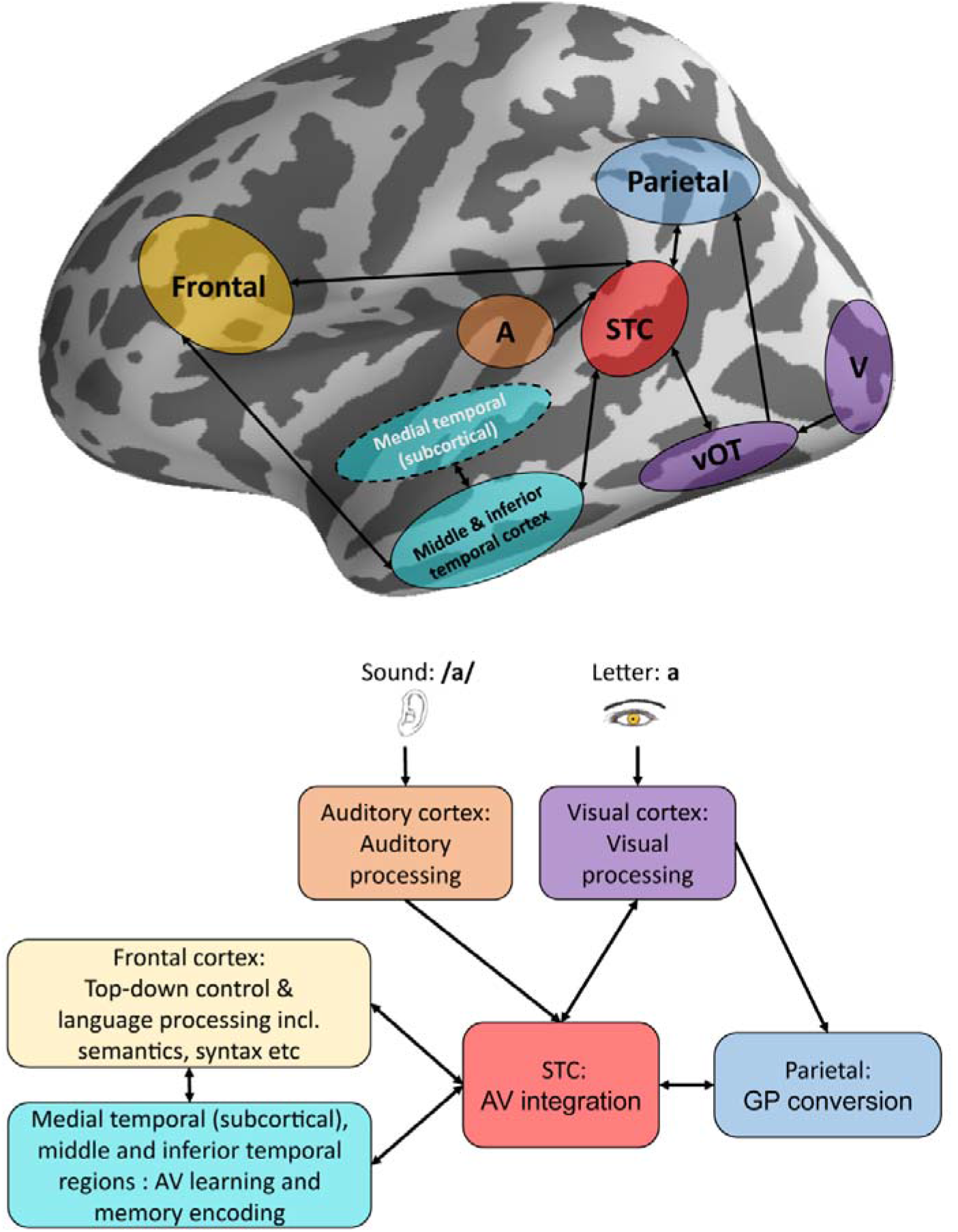
Schematic diagram of the possible network involved in the learning of letters-speech sound associations. A = Auditory cortex, V = Visual cortex, STC = superior temporal cortex, vOT= ventral occipitotemporal cortex, GP = Grapheme-phoneme.

During the initial learning stage, the audiovisual representation is encoded, and short-term memory of the audiovisual objects are stored in the middle and inferior temporal and possibly also in the medial temporal (e.g., hippocampus) regions (Quinn et al. 2017). Frontal regions have been suggested to be involved in many aspects of language-related processes including those related to syntax and semantics (Skeide et al. 2014; Vigliocco 2000), as well as the top-down control mechanism during language learning (Mei et al. 2014; Skeide and Friederici 2016). For example, the frontal regions are involved in the selection of cross-modal features (Hämäläinen et al. 2019; Calvert et al. 2001) to combine and direct attention to the relevant learning cues. In addition, parietal regions also receive visual inputs (of letters) from the occipital regions and might be involved in storing the corresponding phonological representation of the letters by interacting with the multisensory superior temporal cortex during the early stages of learning. As learning progresses, changes have been reported to occur in vOT (Quinn et al. 2017; Madec et al. 2016; Hashimoto and Sakai 2004; Brem et al. 2018; Brem et al. 2010) and dorsal pathway (Taylor et al. 2014; Hashimoto and Sakai 2004; Taylor et al. 2017; Mei et al. 2014; Mei et al. 2015) as well as the STC (Hämäläinen et al. 2019; Karipidis et al. 2018; Karipidis et al. 2017; Madec et al. 2016) for forming optimal cortical representation and automatic processing of the audiovisual objects.

There are several limitations in our study. First, deep brain structures such as the medial temporal system (including the hippocampus) play a crucial role in the learning and memory processes as reported by numerous studies (Jarrard 1993; Brasted et al. 2003; Mayes et al. 2007; Axmacher et al. 2008). One recent fMRI study has also demonstrated that reading-related reorganization could occur at the level of the brainstem and the thalamus (Skeide et al. 2017). MEG might not be optimal to localize the brain activity within these deep brain regions due to the decreased signal-to-noise ratio (SNR) as a function of source depth. However, evidence suggests that hippocampal activities could be captured with MEG (Ruzich et al. 2019; Attal and Schwartz 2013), especially during learning and memory tasks (Taylor et al. 2012; Backus et al. 2016; Shah-Basak et al. 2018). In our study, there seemed to be some activity related to the processing of the learning cues from the deep brain sources. However, due to the limited SNR and spatial resolution in MEG, caution should be taken when interpreting these results, and particularly the localization regarding the medial temporal sources. Another limitation of our study relates to the lack of individual structural magnetic resonance images, which could potentially lead to poorer source localization accuracy and false-positive activation (Supek and Aine 2014).

The current study tracked the learning process in two days; ideally, it would be interesting to track the learning process over a longer period (e.g., one week). In this study, our greatest interest was to investigate the initial stages of learning using a paradigm simulating the situation when children typically learn letters instead of tracking long-term changes in brain activity. Findings from the present study on brain dynamics during letter-speech sound learning could provide new information on potential mechanisms leading to long-term learning outcomes. It is very likely that similar brain networks are recruited for learning letter-speech sound associations in both children and adults since this process utilizes a more general audiovisual object association learning mechanism which is also essential in everyday life. It would be interesting to further investigate the potential difference in behavioral (e.g. learning speed) and brain level differences for adults and children. Learning also involves interaction and communication between different brain regions. Thus brain connectivity would be an interesting approach. The current study has identified important brain regions (hubs) and time windows that could be useful for future studies that use optimal experimental design for connectivity analysis on learning.

In conclusion, in this study, we have successfully captured some of the brain dynamics of learning grapheme-phoneme associations using a well-controlled audiovisual training paradigm. Audiovisual processing showed fast and dynamic changes across different learning stages over two days and was modulated by the effect of overnight memory consolidation during sleep. Newly-learned letters that were associated with specific phonemes showed stronger activation along the dorsal pathway, probably reflecting the grapheme to the phoneme conversion process. We also identified other neural processes, for example, in the middle and inferior temporal cortices, that are important for multisensory learning and cross-modal memory encoding. Letter-speech sound learning deficit has been reported as a key factor for dyslexia in studies using artificial letter training paradigms (Aravena et al. 2013; Aravena et al. 2018; Karipidis et al. 2017; Karipidis et al. 2018). Findings from the present study could provide a better understanding of neural dynamics that underpin grapheme-phoneme learning and could be used to find specific bottlenecks in learning cross-modal associations.

## ACKNOWLEDGEMENTS

We would like to thank Aino Sorsa and Ainomaija Laitinen for their help in data collection. This work has been supported by the European Union projects ChildBrain (Marie Curie Innovative Training Networks, no. 641652), Predictable (Marie Curie Innovative Training Networks, no. 641858) and the Academy of Finland (MultiLeTe #292466). The authors declare no conflicts of interest.

## Notes

https://openneuro.org/datasets/ds002598

### Competing Interest Statement

The authors have declared no competing interest.

### Summary of Updates

Analysis of the congruency effect has been refined. FDR correction applied to the correlation analysis. Figures have been refined.

